# Noise-induced hearing loss delays auditory learning and reduces the influence of acoustic evidence on choice

**DOI:** 10.64898/2026.09.10.750711

**Authors:** Marissa Calvano, Mia Mohr, Willems Mortimer, Samira Alom, Kamala Prerna Annaji Rao Nookala, Todd M. Mowery, Justin D. Yao

**Affiliations:** Department of Head and Neck Surgery & Communication Sciences, Rutgers Robert Wood Johnson Medical School, New Brunswick, NJ, 08901, USA; Brain Health Institute, Rutgers, The State University of New Jersey, Piscataway, NJ, 08854, USA; Department of Psychology – Behavioral and Systems Neuroscience, Rutgers, The State University of New Jersey, Piscataway, NJ, 08854, USA

**Author notes:** Correspondence: Justin D. Yao, Ph.D. Rutgers, The State University of New Jersey Nelson Biological Laboratory D418, 604 Allison Road, Piscataway, NJ 08854. Co-first Author (Marissa Calvano, Mia Mohr).

**Keywords:** noise-induced hearing loss, auditory learning, sensory evidence, Mongolian gerbil

## Abstract

Noise-induced hearing loss (NIHL) degrades auditory sensitivity, but how it affects the learning of sound-guided decisions remains unclear. We trained adult Mongolian gerbils with normal-hearing (NH) or permanent NIHL, induced by a 2-hour exposure to 120 dB SPL broadband noise, to discriminate 4- versus 12-Hz amplitude-modulated (AM) broadband noise presented in a two-alternative forced-choice task. Stimuli were presented at comparable sensation levels across groups. Relative to NH animals, NIHL animals required approximately 2.5 times more sessions and 2.9 times more trials to reach task acquisition criterion. This learning delay was not explained by reduced task engagement, response latency, or auditory brainstem response threshold shift. Video tracking showed that spatial occupancy refined with training in both groups, whereas trial-by-trial movement path trajectories became less variable in NH animals. Per-session and dynamic trial-by-trial logistic regression models revealed that the influence of the task-relevant acoustic stimulus increased with training in both groups but remained consistently lower in NIHL animals, whereas sound-independent choice tendencies were largely unchanged. Together, these results suggest that NIHL delays auditory learning by reducing the influence of acoustic evidence on choice, indicating that hearing loss can alter how sensory experience is used to acquire new behaviors.

## Introduction

Hearing loss may not only impair access to acoustic information but also the learning processes through which auditory experience shapes language and behavior. Acquired hearing loss is associated with changes in auditory perception^1^, but considerably less is known about how it affects the ability to learn new sound-guided behaviors. Human studies suggest that auditory deprivation can impact learning-related processes. For example, children with hearing loss show poorer language and academic outcomes, and hearing-impaired children with cochlear implants show altered sequence learning and memory^2–6^. Adults with age-related hearing loss retain the capacity for auditory perceptual learning, but learning can be slower or reduced compared with normal-hearing (NH) adults^7,8^. Similar differences have been observed during the learning of basic acoustic discriminations^9^. In animal models, hearing loss acquired in adulthood can also impair discrimination of temporally-varying sounds even when differences in sensation level and attentional measures are accounted for^10^. Consistent with these findings, developmental conductive hearing loss in gerbils slows the acquisition of an auditory discrimination task and delays generalization of a learned auditory rule^11^. Together, these studies suggest that hearing loss can alter how experience shapes behavior, but most evidence comes from developmental hearing loss, where auditory deprivation occurs during neurodevelopment, or age-related hearing loss, where sensory decline occurs alongside broader effects of aging. Whether hearing loss acquired after normal auditory development directly alters the subsequent acquisition of a new sound-guided behavior remains unclear.

Noise exposure is a common and preventable cause of acquired sensorineural hearing loss, that can produce permanent damage to cochlear hair cells and auditory-nerve synapses^12–14^. This makes noise-induced hearing loss (NIHL) an experimentally tractable model for determining how a defined auditory insult affects subsequent sound-guided learning. However, compensating for elevated hearing thresholds does not necessarily restore the fidelity of suprathreshold acoustic representations. NIHL can impair frequency selectivity and temporal processing in the auditory periphery^15–17^, and noise exposure produces long-lasting changes in temporal coding and sound representations in auditory cortex^18–20^. Behavioral decision-making deficits after NIHL can also persist when stimuli are presented at comparable sensation levels, indicating that reduced audibility alone does not account for impaired sound-guided behavior^21^. One possibility is that NIHL alters the neural representation of acoustic cues, thereby changing the sensory evidence available to guide behavior^15,17,18,20^. Thus, altered suprathreshold processing after NIHL raises the possibility that auditory learning is impaired not simply by reduced audibility, but by a weaker coupling between acoustic information and sound-guided behavior.

In this study, we tested whether permanent NIHL delays the acquisition of a new sound-guided decision and whether any changes in learning are associated with differences in the influence of acoustic evidence on choice. Adult gerbils with NH and permanent NIHL were trained through acquisition on an amplitude-modulated (AM) rate discrimination task, with stimulus levels adjusted to provide comparable sensation levels across hearing status conditions. We quantified the amount of training required to reach criterion and used video tracking to examine how movement patterns developed across training sessions. To determine what information guided choices throughout training, we fit both per-session and dynamic trial-by-trial logistic regression models that separated the influence of the current acoustic stimulus from sound-independent side bias and previous-choice dependence. We hypothesized that NIHL would delay auditory task acquisition by weakening the process through which acoustic evidence guides choice.

## Results

### Noise exposure produced a permanent elevation of auditory brainstem response thresholds

Adult gerbils (N = 20) were assigned to a normal-hearing group (NH, n = 10; 5 males, 5 fmeales) or a noise-induced hearing loss group (NIHL, n = 10; 7 males, 3 females). NIHL animals received a single 2-h exposure to 120 dB SPL broadband noise (Figure 1A). Before exposure, the groups were indistinguishable as auditory brainstem response (ABR) thresholds did not differ for clicks or for any tested tone frequency (Mann-Whitney U tests, all p ≥ 0.40). Figure 1B shows click-evoked ABRs from one representative NIHL animal before and 14 days after noise exposure. Click thresholds increased from 23 ± 3 dB SPL at baseline to 82 ± 2.5 dB SPL one day after noise exposure, followed by modest recovery to 66 ± 2.2 dB SPL by day 14 (Figure 1C). The day-14 threshold shift was significant in every exposed animal (43 ± 2.6 dB SPL; Wilcoxon signed-rank test, p = 0.002) and remained stable through days 21 and 28 (Friedman test, χ²(2) = 2.00, p = 0.37). The behavioral acquisition phase started 1-2 days after the day-14 ABR recording session, once threshold shifts had stabilized. At day 14, click thresholds in NIHL animals were on average 42 dB SPL higher than in NH animals (66 ± 2.2 vs. 24 ± 3.1 dB SPL; Mann-Whitney U = 0, p < 0.0001) which is equivalent to their own post-14 days threshold shift.

**Figure 1.**
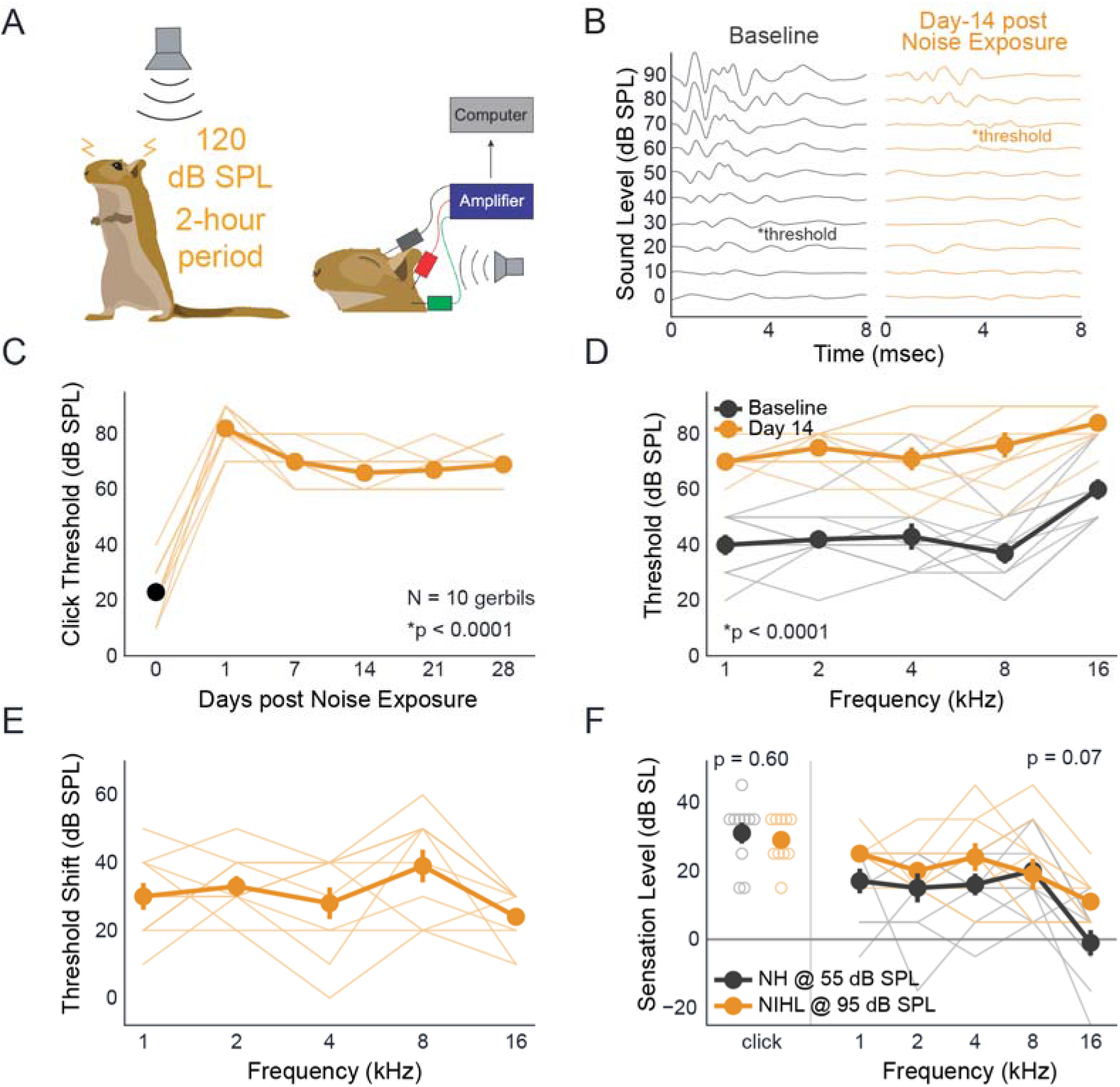
Noise exposure induces permanent broadband hearing loss while preserving comparable task sensation levels. (A) Schematic of noise exposure and ABR recording. NIHL animals received a single 2-hour exposure to 120 dB SPL broadband noise. ABRs were recorded using subdermal electrodes. (B) Representative click-evoked ABR waveforms from one NIHL animal at baseline (grey) and 14 days after exposure (orange). Responses are shown from 0 to 90 dB SPL in 10 dB steps. Asterisks indicate the lowest level at which a reproducible response was identified as threshold. (C) Click ABR thresholds at baseline (day 0, grey) and 1, 7, 14, 21, and 28 days after exposure for every NIHL animal. Thin lines represent individual animals and thick symbols show group mean ± SEM. Click thresholds were significantly elevated by day 14 and remained stable thereafter. (D) ABR thresholds for 1-, 2-, 4-. 8-, and 16-kHz tones at baseline (grey) and 14 days after noise exposure (orange). P-values indicate the hearing status effect and hearing status x frequency interaction, demonstrating a significant broadband elevation in thresholds after noise exposure. (E) Threshold shift (day 14 − baseline) for each tone frequency, demonstrating broadband hearing loss. (F) Sensation level of the behavioral stimulus, calculated as the presentation level minus each animal’s ABR threshold. NH animals were trained at 55 dB SPL and NIHL animals trained at 95 dB SPL. Click sensation level did not differ between groups (p = 0.60), and sensation level across tone frequencies likewise did not significantly differ between groups (p = 0.07). Thin lines and open symbols represent individual animals. Filled symbols represent group mean ± SEM. N = 10 animals per hearing status group.

The hearing sensitivity shift after noise exposure was broadband. Tone thresholds were elevated at each tested frequency 14 days after exposure (Figure 1D-E), with mean shifts of 30 ± 3.9 dB SPL at 1 kHz, 33 ± 3 dB SPL at 2 kHz, 28 ± 4.7 dB SPL at 4 kHz, 39 ± 4.8 dB SPL at 8 kHz, and 24 ± 2.7 dB SPL at 16 kHz. A linear mixed-effects model confirmed a main effect of hearing status across frequency (χ²(1) = 38.4, p < 0.0001) with no hearing status x frequency interaction (χ²(4) = 6.27, p = 0.18). ABR recordings were tested as high as 90 dB SPL, thus several post-noise exposure tone thresholds were reported at that ceiling value and the reported tone shifts therefore underestimate the loss in those animals.

Task stimuli were presented at comparable sensation levels across hearing status conditions. Differences in task acquisition rate after noise exposure could be potentially confounded if the task stimulus were simply less audible to NIHL animals. We therefore presented the AM stimuli at 55 dB SPL for NH animals and 95 dB SPL for NIHL animals and expressed each level relative to an animal’s own ABR threshold (Figure 1F). For clicks, the stimulus closest in bandwidth to the broadband AM task stimulus, sensation level did not differ between groups (NH, 31 dB sensation level; NIHL, 29 dB sensation level; difference = −2 dB, 95% CI [−9.9, +5.9], t(18) = −0.53, p = 0.60). Across the five tone frequencies, sensation level was 6.4 dB higher in the NIHL group, a difference that did not reach significance (95% CI [−0.5, +13.3], t(18) = 1.95, p = 0.07). Thus, the task-relevant stimulus was not presented at a lower sensation level after noise exposure compared to NH conditions.

### Noise-induced hearing loss delays acquisition of an amplitude-modulated rate discrimination task

Following procedural shaping, gerbils were trained to discriminate 4-versus 12-Hz amplitude-modulated (AM) broadband noise in a single-interval, two-alternative forced-choice task (Figure 2A). Animals self-initiated each trial at a central nose poke and reported the AM rate by approaching the left food trough for 4-Hz or the right food trough for 12-Hz. Task acquisition was defined as ≥80% correct on both AM rates within a single session. Representative learning curves are shown as a function of training session (Figure 2B-E) and cumulative trials (Figure 2F-I). Learning curves for every animal are shown in Supplementary Figures S1 and S2.

**Figure 2.**
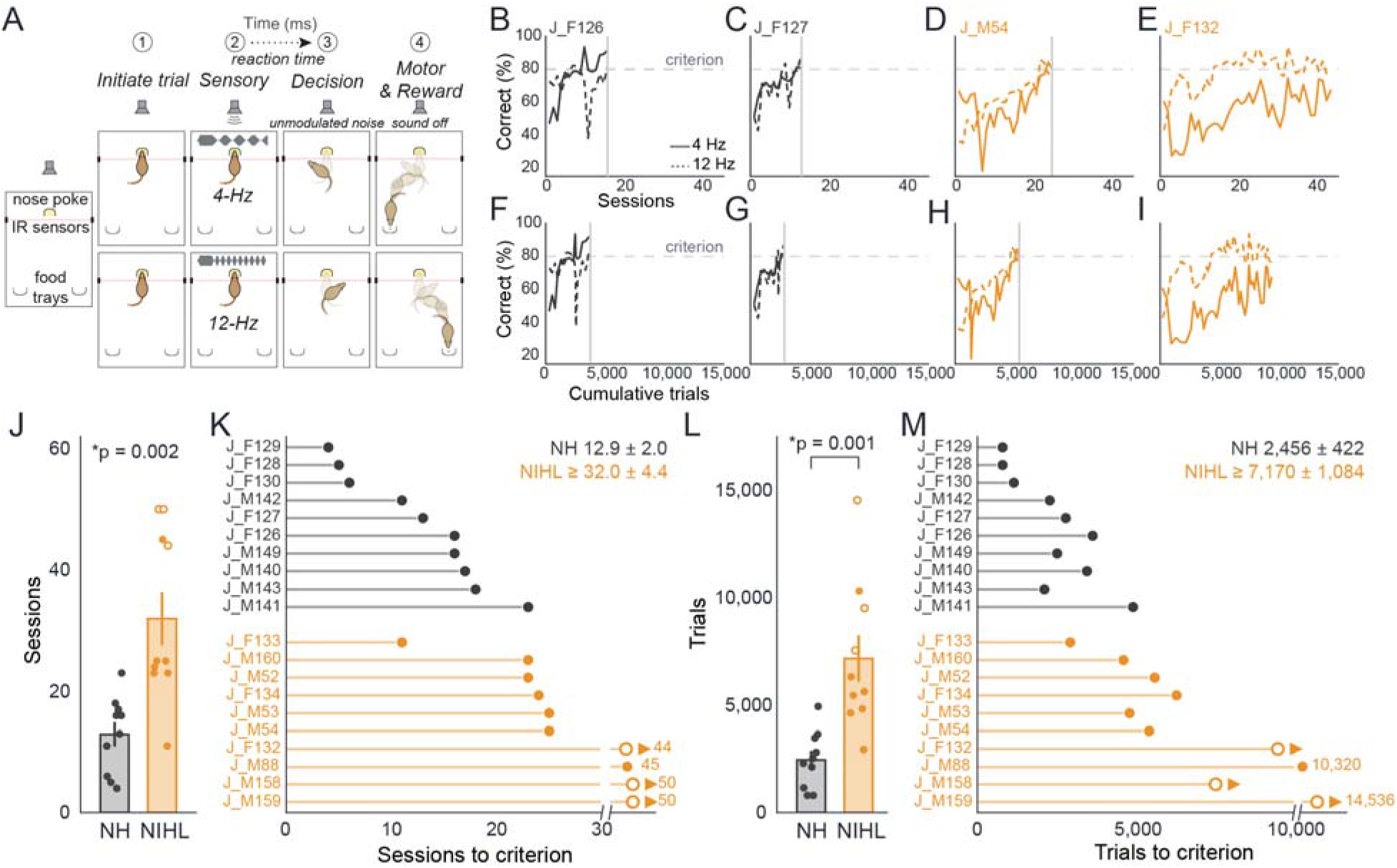
Noise-induced hearing loss delays acquisition of the auditory discrimination task. (A) Schematic of the behavioral task. (1) The animal initiates a trial at the central nose poke, breaking an infrared beam. (2) This triggers presentation of an AM stimulus presented at either 4-or 12-Hz. (3) The animal makes its decision by approaching to left food trough for 4-Hz trials or right food trough for 12-Hz trials, and the sound transitions to unmodulated noise. (4) Correct responses are rewarded with 20 mg pellets. (B–E) Percent correct for each AM rate across sessions (solid, 4-Hz; dashed, 12-Hz) for two representative NH animals and two NIHL animals. The horizontal dashed line marks the 80% acquisition criterion and the vertical grey line indicates the session at which criterion was first met on both AM rates. Animal J_F132 was discontinued without reaching the criterion. (F–I) The same representative animals plotted as function of cumulative trials. (J) The number of sessions required to reach criterion for NH and NIHL animals. Bars are the group mean ± SEM and points are individual animals. Open symbols are the three NIHL animals discontinued without reaching criterion. The p-value indicates the Mann-Whitney comparison between hearing status groups. (K) The number of sessions to criterion for each individual animal. Open symbols with arrows indicate discontinued animals, for which the plotted value represents the session at which training stopped and therefore a lower bound. The axis is broken at 30 sessions and the off-scale values are printed. (L, M) The same as J-K, for cumulative trials. The p-value in L indicates the Mann-Whitney comparison between hearing status groups for trials to criterion. N = 10 per hearing status group.

NIHL animals required substantially more training to reach task acquisition criterion. NH animals reached criterion in 12.9 ± 2 sessions, whereas NIHL animals required at least 32 ± 4.4 sessions (Mann-Whitney U = 7.5, p = 0.002; Figure 2J-K). The same effect was present when training was expressed as cumulative trials across all sessions as NH animals required 2,456 ± 422 trials versus at least 7,170 ± 1,084 trials for NIHL animals (Mann-Whitney U = 5, p = 0.001; Figure 2L-M). Thus, NIHL animals required approximately 2.5 times as many sessions and 2.9 times as many trials to acquire the task.

All 10 NH animals reached criterion, compared with 7 of 10 NIHL animals. The remaining three NIHL animals were discontinued at sessions 44 or 50 after prolonged periods without improvement. Their plotted session and trial counts are therefore lower bounds rather than completion times, which makes the NIHL group averages and the rank-based comparisons conservative estimates. The group difference also did not depend on placing the acquisition criterion specifically at 80% as it remained significantly different at 70% and 75% criteria for both sessions and trials (Mann-Whitney U tests, all p ≤ 0.002).

Exploratory analyses were performed to assess potential sex differences in task acquisition within each hearing status group. Among NH animals, there were no significant differences between females and males in the number of sessions (females = 8.8 ± 5.4; males = 17 ± 4.3 sessions; Mann-Whitney U = 2.5, p = 0.06) or cumulative trials (females = 1,842 ± 1,308, males = 3,070 ± 1,164 trials; Mann-Whitney U = 7, p = 0.31) required to reach task acquisition criterion. Within the NIHL group, 2 of 3 females and 5 of 7 males reached task acquisition criterion. Exploratory time-to-criterion analyses did not reveal a significant difference between sexes when task acquisition was compared across either training sessions (log-rank test, p = 0.51) or cumulative trials (log-rang test, p = 0.85). These comparisons among NIHL animals were limited by the small number of females (n=3), but the available data did not indicate a significant sex-dependent difference.

If slower learning were a direct consequence of reduced audibility, NIHL animals with the largest threshold shifts should have required the most training. However, we did not observe this relationship. Across the primary comparisons of click threshold shift and mean tone threshold shift, each against sessions and trials to criterion, we found no significant relationship (Spearman rank correlations, n = 10; Holm-corrected p-values = 0.14, 0.65, 0.93, and 1). Absolute day-14 thresholds and threshold shifts at individual tone frequencies were likewise unrelated to task acquisition (all p ≥ 0.12). Thus, these results suggest the magnitude of NIHL does not predict task acquisition.

### Slower task acquisition in noise-induced hearing loss animals was not explained by reduced task engagement

NIHL animals remained comparatively engaged during task performance across training compared to NH animals. Across the full training record, NIHL animals completed more trials per session than NH animals (NIHL = 227 ± 13; NH = 190 ± 11; Mann-Whitney U = 22, p = 0.04), and the groups were indistinguishable when compared over the first five or first ten sessions (both p ≥ 0.27). The percentage of failure trials with no choice response did not differ between groups (NIHL = 1.57 ± 0.47%, NH = 1.71 ± 0.22%; Mann-Whitney U = 67,p = 0.21), as was the percentage of trials aborted (NIHL = 0.54 ± 0.24%; NH = 0.52 ± 0.21%; Mann-Whitney U = 49, p = 0.97). In addition, NIHL animals were also comparable to NH animals in the speed of making their choices. Median response latency did not differ between groups across training (NIHL = 1,232 ± 168 ms; NH = 1,147 ± 143 ms; Mann-Whitney U = 41, p = 0.52), even when the analysis was restricted during the first five sessions (p = 0.24), or to only correct trials (p = 0.39). Thus, across the number of completed trials, failures to respond, aborted trials, and response latency, NIHL animals remained at least as engaged with the task as NH animals. This suggests reduced task participation therefore does not account for the longer task acquisition duration.

### Movement patterns refine during learning but remain less stereotyped after NIHL

Video tracking of head position was available for 14 animals (NH, n = 8; NIHL, n = 6). For each session, we quantified the proportion of the test cage arena visited by the animal’s head. Representative occupancy maps show that the broad, diffuse coverage present early in training became concentrated into more defined routes between the nose poke and food troughs by task acquisition in both groups (Figure 3C, E). There was a strong negative relationship between area occupied across sessions in the representative NH animal (Spearman rank correlation, ρ = −0.90, p = 0.04; Figure 3D) and NIHL animal (Spearman rank correlation, ρ = −0.84, p < 0.001; Figure 3F). This same refinement occurred across animals as the percentage of arena occupied declined over normalized training progress in every tracked animal in both groups (NH: Spearman’s ρ = −0.70 ± 0.06, Wilcoxon signed-rank p = 0.008; NIHL: Spearman’s ρ = −0.59 ± 0.14, p = 0.03; Figure 3G-H). Comparing the first and second half of training sessions produced a similar pattern where the percentage of arena occupied significantly decreased in NH animals (Wilcoxon signed-rank, p = 0.02) and showed a decreasing trend in NIHL animals (Wilcoxon signed-rank, p = 0.06) (Figure 3I). No between-group difference was detected in percentage arena occupied at any quarter of training (all p ≥ 0.49). These results suggest NIHL delayed acquisition without preventing the general spatial refinement that accompanies task experience and learning.

**Figure 3.**
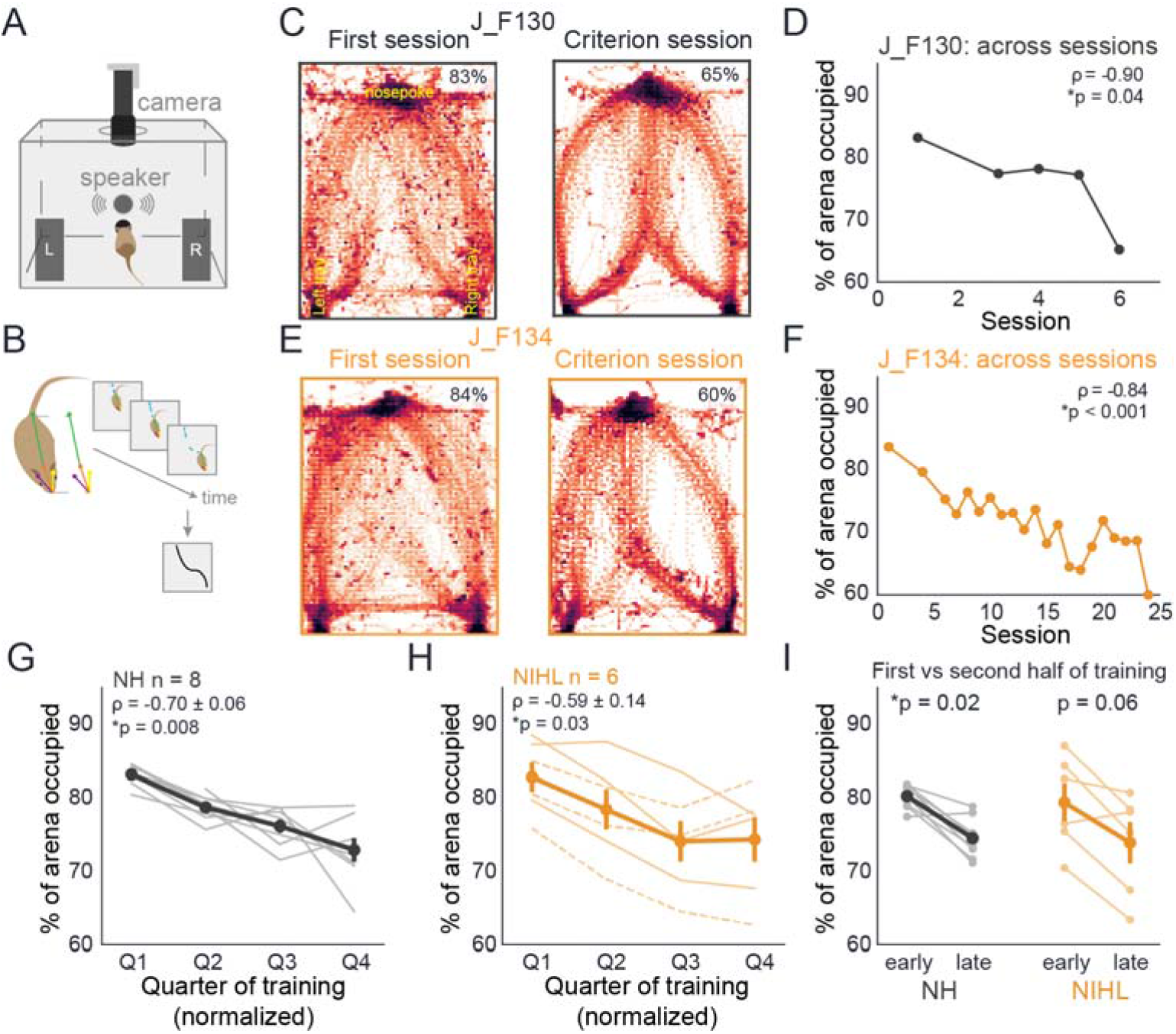
Spatial exploration decreases across training in both normal-hearing and noise-induced hearing loss animals. (A) Schematic of the behavioral arena and overhead camera used for video tracking. (B) Head position was tracked frame-by-frame with SLEAP^52^ to quantify movement through the arena across training. (C) Representative head-occupancy maps for an NH animal on its first session and on the session at which it reached task acquisition criterion. Color represents dwell density where darker regions represent the animal spent more time there. Values indicate the percentage of the arena occupied. (D) Percentage of arena occupied across training sessions for the same NIH animal. Spearman’s ρ and corresponding p-value indicate the relationship between training session and percentage of arena occupied. (E, F) Corresponding occupancy maps and session-by-session percentage of arena occupied for a representative NIHL animal. Spearman’s ρ and corresponding p-value indicate the relationship between training session and area occupied. (G-H) Percentage of arena occupied across normalized quarters of training in NH and NIHL animals. Thin lines represent individual animals and thick lines show group mean ± SEM. Mean Spearman’s ρ represents the average within-animal relationship between training session and percentage of arena occupied, and p-values indicate Wilcoxon signed-rank tests of the per-animal correlation coefficients against zero. Dashed lines indicate NIHL animals discontinued without reaching criterion. (I) Comparison of the first and second halves of training. P-values indicate within-group Wilcoxon signed-rank comparisons between training progression halves.

A reduction in the total area visited does not indicate whether an animal takes the same route to and from the nose poke to the food troughs from trial-to-trial. We therefore quantified movement trajectory path spread, defined as the mean distance of individual correct-trial movement trajectories from each session’s mean route to each food trough (Figure 4A). In the representative NH animal, movement trajectories became progressively more stereotyped as path spread fell from 28 to 17 pixels across training session (Spearman ρ = −0.90, p = 0.04; Figure 4B-C). The representative NIHL animal reached task acquisition criterion without a comparable monotonic decrease in trajectory path spread (Spearman ρ = 0.34, p = 0.13; Figure 4D-E). Across animals, path spread declined in every NH animal across normalized training sessions (mean Spearman ρ = −0.59 ± 0.10, Wilcoxon signed-rank p = 0.008; Figure 4F) but not in the NIHL group, in which five of six animals showed a flat or increasing trend (Spearman ρ = 0.34 ± 0.19, p = 0.22; Figure 4G). NH animals displayed a significant decrease from the first to second half of training (Wilcoxon signed-rank p = 0.02), whereas no change occurred for NIHL animals (Wilcoxon signed-rank p = 0.16) (Figure 4H). These data suggest that NH animals increasingly converged on a repeatable action path across training, whereas this shift in trajectory consistency did not occur in NIHL animals.

**Figure 4.**
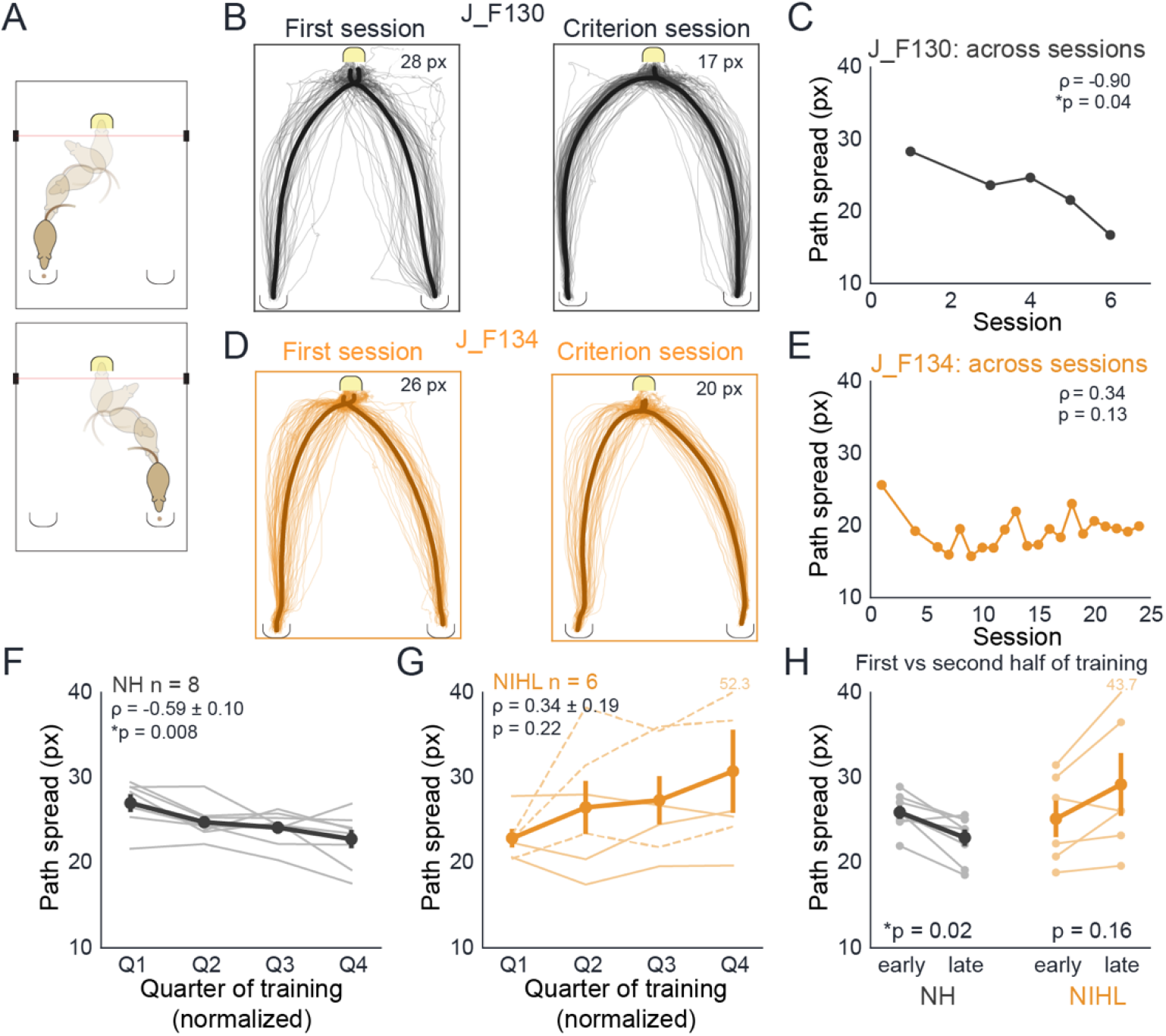
Movement trajectories become more consistent with training in normal-hearing animals but in noise-induced hearing loss animals. (A) Schematic illustrating extraction of head trajectories from the central nose poke to the left or right food trough on correct trials. (B) Trial-by-trial trajectories from a representative NH animal during the first session and criterion session. Thin lines indicate individual trajectories and thick lines indicate the mean route to each food trough. Path spread, defined as the mean distance of individual trajectories from the session mean route, decreased from 28 to 17 pixels. (C) Path spread across training sessions for the same NH animal. Spearman’s ρ and corresponding p-value indicate the relationship between training session and path spread. (D-E) Corresponding trajectories and session-by-session path spread for the representative NIHL animal. Spearman’s ρ and corresponding p-value indicate the relationship between training session and path spread. (F-G) Path spread across normalized quarters of training for NH and NIHL animals. Thin lines represent individual animals and thick lines show group mean ± SEM. Mean Spearman’s ρ represents the average within-animal relationship between training session and path spread, and p-values indicate Wilcoxon signed-rank tests of the per-animal correlation coefficients against zero. Dashed lines indicate NIHL animals discontinued without reaching criterion. (H) Comparison of the first and second halves of training. P-values indicate within-group Wilcoxon signed-rank comparisons between training halves.

### Noise-induced hearing loss reduces the influence of acoustic evidence on choice throughout training

There are a number of factors that could influence a sound-guided decision during task performance. Percent correct describes whether a choice was rewarded, but not what information drove that choice. We therefore fit each training session with a logistic regression model containing three terms: the presented sound, a standing side bias, and the animal’s previous choice. The sound weight measures how strongly the AM stimulus influenced the current choice, and the side-bias and previous choice weights capture sound-independent choice tendencies. Fits were obtained for 437 sessions from all 20 animals (Figure 5). The influence of the sound increased with training in every animal in both groups (NH: Spearman’s ρ = 0.83 ± 0.05, p = 0.002; NIHL: Spearman’s ρ = 0.76 ± 0.03, p = 0.002; Figure 5A-B). However, when animals were compared at matched normalized training progress, the sound weight remained lower among NIHL animals across training (mixed-model group term, χ^2^(1) = 17.8, p < 0.0001; Figure 5C). The difference was already present during the first quarter of training and persisted through the final quarter, but the rate of increase across training did not differ between groups (group x quarter interaction, χ^2^(3) = 4.36, p = 0.23). This suggests both groups increasingly relied on the acoustic stimulus as training progressed, but NIHL animals did so from a consistently lower level.

**Figure 5.**
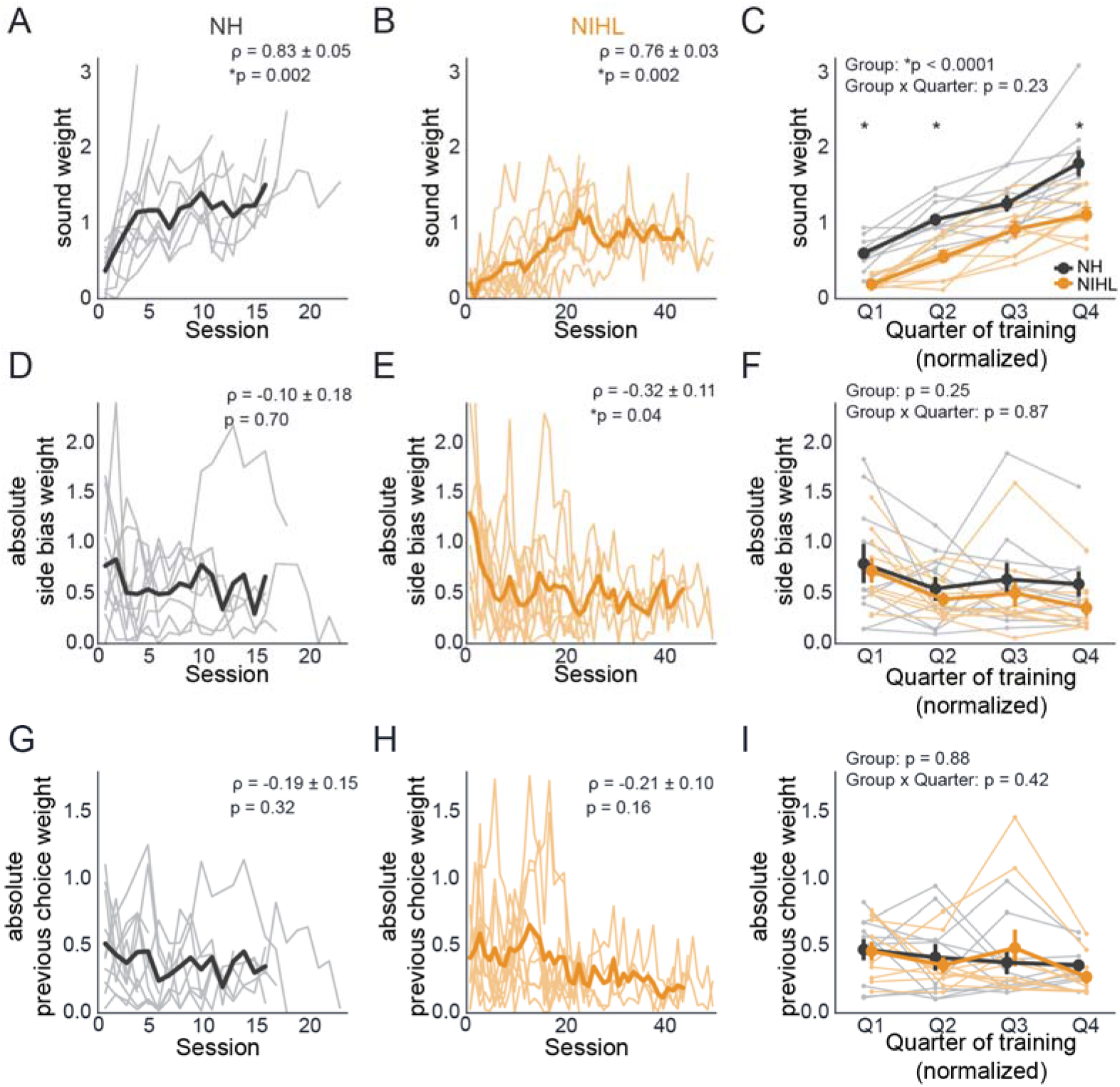
The influence of sound on choice increases across training but remains lower in noise-induced hearing loss animals. Each training was fit separately with a logistic regression model incorporating the current sound, side bias, and previous choice. (A-B) Sound weight across training sessions for NH (A) and NIHL (B) animals. Thin lines represent individual animals and thick lines represent the group mean. Mean Spearman’s ρ and corresponding p-values demonstrate sound weight increased significantly across sessions in both groups. (C) Sound weight across normalized quarters of training. Thick lines and symbols show group mean ± SEM and thin lines show individual animals. P-values represent hearing status group effect and group x training quarter interaction effects, demonstrating sound weight was significantly lower in NIHL animals across training, while the progression across training did not significantly differ between groups. Asterisks indicate significant quarter-wise NH versus NIHL comparisons after Holm correction. (D-E) Absolute side-bias weight across sessions for NH (D) and NIHL (E) animals. Mean Spearman’s ρ and p-value are based on correlations within each group across training sessions. (F) Absolute side-bias weight across normalized training progress. P-values represent statistical comparisons between groups and group x quarter training interaction. (G-H) Absolute previous-choice weight across sessions for NH (G) and NIHL (H) animals. Mean Spearman’s ρ and p-value are based on correlations within each group across training sessions. (I) Absolute previous-choice weight across normalized training progress. P-values represent statistical comparisons between groups and group x quarter training interaction.

Normalizing the progress of learning across sessions may obscure how much training was required to produce the change in the influence of the sound weight. One quarter of training corresponded to substantially more training for NIHL animals (7.8 ± 1.1 sessions) than for NH animals (3.1 ± 0.5 sessions) (Mann-Whitney U = 8, p = 0.001), and thus similar changes across normalized training progress occurred over considerably more actual training in NIHL animals. Accordingly, although the sound weight rose similarly for both groups as a function of normalized training progress, it increased significantly more slowly per training session in NIHL animals (NH: 0.23 ± 0.08; NIHL: 0.06 ± 0.02 weight units/session; Mann-Whitney U = 81, p = 0.02). This suggests NIHL therefore did not prevent animals from learning to use the sound to guide choices, but reduced the influence of acoustic evidence on choice and stretched that learning over substantially more task sessions.

When examining sound-independent choice tendencies, we found no significant relationship between side-bias weight and training session in NH animals (mean Spearman’s ρ = −0.10 ± 0.18, p = 0.70; Figure 5D), whereas side-bias weight significantly decreased across sessions in NIHL animals (Spearman’s ρ = −0.32 ± 0.11, p = 0.04; Figure 5E). This suggests that NIHL became less reliant on an initial fixed side preference as training progressed, whereas NH animals did not show a consistent monotonic change in side bias across sessions possibly due to starting with relative little side bias. Importantly, the overall side-bias trajectories did not differ between groups (mixed-model group, χ²(1) = 1.31, p = 0.25) or during training progression (group x quarter interaction, χ²(3) = 0.72, p = 0.87; Figure 5F). Previous choice weight did not change significantly across training in either group (NH: mean Spearman’s ρ = −0.19 ± 0.15, p = 0.32; NIHL: Spearman’s ρ = −0.21 ± 0.10, p = 0.16; Figure 5G-H). Consistent with this, previous choice weight did not differ between groups in overall level (mixed-model group, χ²(1) = 0.02, p = 0.88) or in its progression across training (interaction, χ²(3) = 2.83, p = 0.42; Figure 5I). Thus, NIHL animals were not compensating for a weaker influence of the sound by relying more strongly on a fixed side preference or on their preceding choice.

### Trial-by-trial logistic regression modeling further reveals weaker stimulus-choice coupling after noise-induced hearing loss

The per-session analysis treats each session as a single unit. To resolve how choice strategy developed within and across sessions, we fit the same three terms (i.e., sound, previous choice, and side bias) with a dynamic logistic regression model in which each weight was allowed to evolve from trial-to-trial as a Gaussian random walk^22^ (Figure 6A). One model was fit to each animal’s complete training record, comprising 94,871 choice trials across 449 sessions. The dynamic estimate closely tracked the corresponding per-session fits while resolving changes within sessions (Figure 6B). Complete trial-by-trial fits for every animal are shown in Supplementary Figure S3.

**Figure 6.**
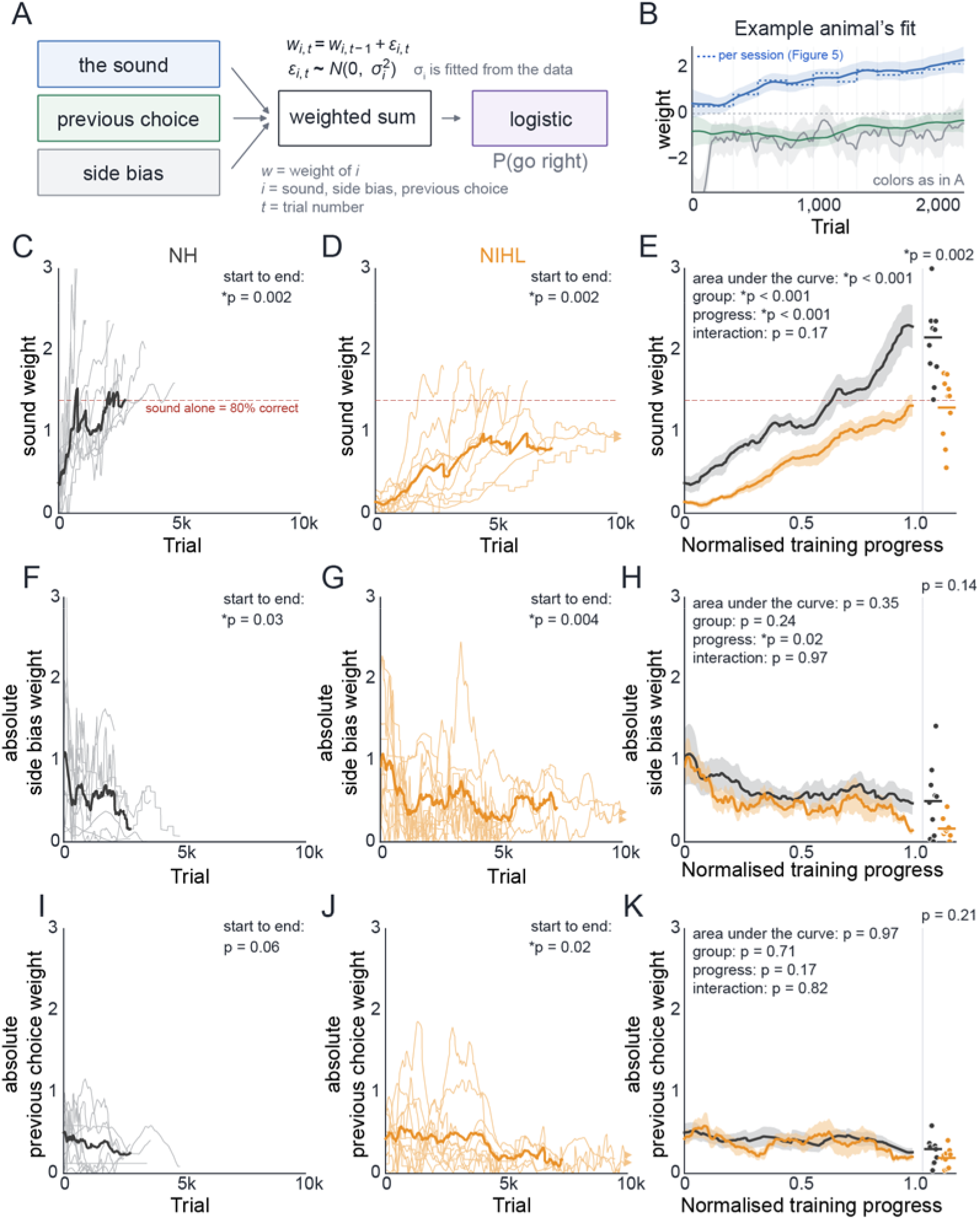
Trial-by-trial modeling reveals persistently weaker stimulus-choice coupling in noise-induced hearing loss animals. (A) Schematic of the dynamic logistic regression model. The current sound, previous choice, and side bias are combined as weighted sum and transformed through a logistic function to estimate the probability of choosing the right response. Unlike the per-session model in Figure 5, each weight is allowed to evolve from trial-to-trial according to a random-walk process, with the step size estimated from the data, as implemented in PsyTrack^22^. (B) Example trial-by-trial fit from one animal. Solid lines show the dynamic model weights with shaded ± 2 posterior SD, and the dotted blue line shows the corresponding per-session sound weight from Figure 5 for comparison. (C-D) Trial-by-trial sound weight across training for NH (C) and (D) animals. Thin lines represent individual animals and thick lines represent the group mean. P-values indicate within-group comparisons between the beginning and end of training. The dashed red line indicates the sound weight at which the sound alone would support 80% correct performance. (E) Sound weight across normalized training progress. Thick lines show group mean ± SEM and individual end-of-training value are shown at on the right. P-values indicate the area-under-the-curve comparison between hearing status groups, as well as mixed-model group, training-progress, and group x progress interaction effects. (F-G) Absolute side-bias weight across trials for NH (F) and NIHL (G) animals. P-values indicate within-group comparisons between the beginning and end of training. (H) Absolute side-bias weight across normalized training progress. P-values indicate the area-under-the curve comparison between groups and mixed-model group, progress, and group x progress interaction effects. (I-J) Absolute previous-choice weight across trials for NH (I) and NIHL (J) animals. P-values indicate within-group comparisons between the beginning and end of training animals. (K) Absolute previous-choice weight across normalized training progress. P-values indicate the area-under-the-curve comparison between groups and mixed-model group, progress, and group x progress interaction effects.

The sound weight significantly increased from the beginning to the end of training in every animal in both groups (NH: 0.45 to 2.16; NIHL: 0.13 to 1.3; Wilcoxon signed-rank p = 0.002; Figure 6C-D), yet it remained lower in NIHL animals. The groups differed during the first 200 trials (NH: 0.45 ± 0.08; NIHL: 0.13 ± 0.02; Mann-Whitney U = 81, p = 0.02), at the end of training (NH: 2.16 ± 0.20; NIHL 1.3 ± 0.13; Mann-Whitney U = 90, p = 0.002), and across the full trajectory (area under the normalized-progress curve; NH: 1.18 ± 0.10, NIHL: 0.65 ± 0.05; Mann-Whitney U = 96, p < 0.001; Figure 6E). A mixed model confirmed a strong effect of training progress (χ^2^(3) = 81.7, p < 0.001) and hearing status (χ²(1) = 16.5, p < 0.001), with no significant interaction (χ^2^(3) = 5, p = 0.17). These results indicate that the sound became increasingly influential with training in both groups, but it remained consistently less influential in NIHL animals. This suggests reduced stimulus-choice coupling after NIHL, with acoustic evidence exerting less influence on choice throughout training.

The sound-independent terms did not differ between hearing status groups. Absolute side bias weight declined with training in both groups (Wilcoxon signed-rank NH p = 0.03, NIHL p = 0.004; mixed model progress term, χ^2^(3) = 10.3, p = 0.02; Figure 6F-H). However, absolute side bias weight but did not differ between groups across the full trajectory (area under the normalized-progress curve, Mann-Whitney U = 63, p = 0.35; mixed model group term, χ^2^(1) = 1.39, p = 0.24) and the pattern of change across training was similar between groups (mixed model group x progress interaction, χ^2^(3) = 0.27, p = 0.97; Figure 6H). Absolute previous choice weight decreased from the start to the end of training in NIHL animals (Wilcoxon signed-rank test, p = 0.02; Figure 6J). NH animals showed a similar decreasing trend but it did not reach significance (Wilcoxon signed-rank test, p = 0.06; Figure 6I). Across normalized training progress, there was no significant overall effect of training progress (mixed-model progress effect,, χ^2^(3) = 5, p = 0.17; Figure 6K), no group difference (χ^2^(1) = 0.14, p = 0.71), and no group x progress interaction (χ^2^ (3) = 0.92, p = 0.82). The area under the normalized-progress curve, likewise, did not differ between groups (Mann-Whitney U = 51, p = 0.97). Together, the per-session and trial-by-trial analyses identify the same learning deficit where after NIHL, acoustic evidence exerted less influence on choice throughout task acquisition, while sound-independent choice tendencies were largely unchanged.

## Discussion

NIHL substantially delayed acquisition of an auditory discrimination task despite comparable stimulus audibility and preserved behavioral engagement compared to NH control animals. NIHL animals required more sessions and trials to reach learning criterion. This delay was accompanied by a persistent reduction in the influence of acoustic stimulus on choice. Although the influence of task-relevant sound cues increased across training in both groups, it remained consistently lower for NIHL animals, whereas side bias and previous-choice dependence were largely unchanged. Spatial exploration within the test arena also became more restricted with training in both groups, but movement trajectories became increasingly consistent in NH animals. Together, these findings extend our previous work showing impaired auditory decision-making after permanent NIHL^21^ by demonstrating that hearing loss also disrupts the acquisition of sound-guided behavior, particularly the process by which acoustic evidence gains reliable guidance over choice decisions.

The prolonged acquisition after NIHL is unlikely to reflect a simple failure to hear the task-relevant stimulus or reduced engagement during training. Behavioral stimuli were presented at comparable sensation levels between hearing status groups, and the magnitude of ABR threshold shift did not predict the number of sessions or trials required to reach learning criterion. Relative to NH control animals, NIHL animals also completed comparable or greater numbers of trials per session, with no increase in failure or abort trials and no difference in response latency. These findings argue against reduced task-related audibility or task participation as sufficient explanations for the learning deficit we found. Consistent with this interpretation, developmental conductive hearing loss in gerbils was previously shown to delay acquisition of an AM-rate discrimination despite stimuli being presented at comparable sensation levels^11^. However, increasing task-related stimulus sound levels for NIHL animals to balance sensation level does not necessarily restore the fidelity of suprathreshold auditory representations. Adult-onset hearing loss in gerbils similarly impairs temporal auditory perception despite compensation for sensation level, with performance consistent with increased internal sensory noise^10,23^. Noise exposure can produce peripheral neural damage and functional deficits^13,24^ that are not captured by threshold sensitivity alone, and our previous work similarly found impaired auditory decision-making after NIHL despite testing animals at comparable sensation levels to pre-NIHL conditions^21^. Thus, the slower task acquisition observed here may reflect a deficit in how suprathreshold acoustic information is represented or used to guide behavior rather than whether the stimulus can simply be detected.

Our per-session choice-modeling analysis and PsyTrack-based trial-by-trial dynamic modeling^22^ revealed that NIHL reduced the influence of acoustic evidence on behavior throughout learning. NIHL animals learned to use the acoustic stimulus to guide behavior, but its influence on choice remained consistently weaker throughout training compared to NH animals. This was evident early in training and persisted through the end of task acquisition. Thus, NIHL did not prevent animals from learning to use the sound, but weakened the degree to which stimulus identity controlled their choices. Stimulus-independent strategies, including choice bias and previous-choice dependence, can evolve across auditory category learning and contribute to individual learning trajectories^25^. Importantly, this reduction was not accompanied by greater reliance on sound-independent strategies. Side bias and previous-choice dependence were largely similar between groups, indicating that NIHL animals were not simply compensating for weaker acoustic information by adopting a fixed response bias or relying more strongly on recent choice history. Together, these findings suggest that the prolonged task acquisition after NIHL reflects reduced stimulus-choice coupling, such that more training may be required for acoustic evidence to gain reliable influence over behavior.

The reduced influence of task-relevant acoustic cues on choice could arise from several non-mutually exclusive mechanisms. First, NIHL may degrade the fidelity of suprathreshold auditory representations, such that the sensory evidence available on each trial is noisier or less discriminable even when stimuli are presented at comparable sensation levels. Noise-induced peripheral damage can substantially alter temporal-envelope coding^26^, while compensatory central gain may restore some measures of sound responsiveness without fully restoring the representation of temporally-modulated signals^27^. At the cortical level, developmental hearing loss can degrade population encoding of AM cues, with perceptual deficits associated with greater trial-to-trial variability in auditory cortical responses^23^. Moreover, cochlear neural degeneration can increase internal noise within auditory cortex and disrupt perceptual performance despite relatively preserved sound detection^20^. Second, acoustic information may remain sufficiently represented within auditory cortex but is transmitted to or read out less effectively by downstream circuits. Animal models and human imaging studies have demonstrated hearing loss can alter functional interactions between auditory and non-auditory cortical regions^28–32^, while parietal, frontal, and corticostriatal circuits are known to transform and accumulate sensory evidence during perceptual decisions^33–37^. Finally, NIHL may interfere with the learning-related plasticity through which behaviorally-relevant acoustic representations become increasingly linked to choices and actions. Auditory cortical representations are dynamically modified by task engagement and learning^38–44^, including for temporally-modulated sounds, where behaviorally-relevant categorical representations emerge within deep-layer auditory cortical output neurons^45^. Consistent with this possibility, developmental hearing loss in gerbils has been shown to slow auditory task acquisition and delay generalization of learned auditory associations^11,46^, as well as impair perceptual learning^47^. Additionally, auditory learning is accompanied by plastic changes within downstream corticostriatal circuits that are altered following developmental auditory deprivation^48^. These findings raise the possibility that NIHL slows or limits this experience-dependent transformation. These mechanisms are not mutually-exclusive, and the present behavioral models cannot determine whether the reduced sound weight originates from altered sensory coding, impaired downstream readout, or disrupted learning-related plasticity.

Our video-recorded movement-tracking data further suggest that NIHL alters the refinement of learned behavior. Spatial exploration decreased across training in both groups, indicating that animals developed a progressively more restricted spatial strategy as they gained experience with the task. However, trial-to-trial movement trajectories became increasingly consistent in NH animals, but not in NIHL animals. Reduction in movement variability is a common feature of skill acquisition and can reflect the progressive stabilization of task-relevant behavioral patterns with practice^49,50^. Thus, the persistent path trajectory variability in NIHL animals may reflect less complete stabilization of the sensorimotor sequence linking stimulus sampling to choice. One possibility is that this is a consequence of weaker stimulus-choice coupling, such that if acoustic evidence provides less reliable influence over choice, the behavioral sequence that follows stimulus presentation may also remain more variable across trials. Alternatively, greater path trajectory variability could represent continued behavioral exploration or a compensatory strategy when sensory evidence is uncertain, as variability can facilitate exploration during learning rather than simply reflecting poor motor control^51^. The dissociation between reduced spatial exploration and persistent path trajectory variability suggests that NIHL does not broadly impair behavioral refinement, but may selectively limit the emergence of a stable sound-guided behavioral strategy.

Together, our findings support a framework in which hearing loss disrupts not only auditory sensitivity, but also the process by which acoustic evidence gains reliable guidance over behavior. By examining task acquisition after adult-onset permanent NIHL, we show that animals retained the capacity to learn from task-relevant sounds, but required substantially more experience than NH animals for those sounds to exert reliable influence over choice. This deficit occurred despite comparable stimulus sensation levels and preserved task engagement, and was accompanied by reduced stabilization of movement trajectories during learning. The present behavioral analyses cannot determine whether these effects originate from degraded auditory representations, altered communication with downstream decision circuits, or impaired learning-related plasticity. Determining how these processes evolve across acquisition will require future experiments of longitudinal measures of sensory and decision-related neural activity. More broadly, these findings suggest that the consequences of hearing loss extend beyond reduced audibility to include changes in how sensory evidence is learned, weighted, and ultimately used to guide behavior.

## Methods

### Experimental design

Adult Mongolian gerbils (*Meriones unguiculatus*; N = 20; 8 females, 12 males) were assigned to a normal-hearing group (NH, n = 10; 5 females, 5 males) or a noise-induced hearing loss group (NIHL, n = 10; 3 females, 7 males). All animals underwent baseline auditory brainstem response (ABR) recording. NIHL animals then received a single noise exposure followed by repeated post-exposure ABR recordings, whereas NH animals received no exposure. Behavioral acquisition in NIHL animals began after hearing thresholds had stabilized post-2 weeks after noise exposure. Both groups were subsequently trained on the same auditory discrimination task. The primary behavioral endpoint was the amount of training required to reach the acquisition criterion, expressed as both sessions and cumulative trials.

### Noise exposure

NIHL was induced with the same loud-noise exposure paradigm used previously in our laboratory^21^. Awake gerbils received a single 2-h exposure to 120 dB SPL broadband noise. ABRs were recorded before exposure and at 1, 7, 14, 21, and 28 days afterward. Behavioral acquisition for NIHL animals began 1-2 days after the day-14 ABR, when threshold shifts had stabilized (see Figure 1C-E).

### Single-interval alternative forced choice auditory discrimination task

Behavioral procedures were adapted from our previously described gerbil auditory decision-making paradigm^21^. Animals were maintained on controlled food access during behavioral training. Testing was performed in a behavioral arena housed within a sound-attenuating cubicle or sound-attenuation booth (Med Associates). A central nose poke port was positioned opposite two food troughs, with infrared sensors used to detect trial initiation, departure from the nose poke region, and entry into each trough (Figure 2A). Stimulus presentation, reward delivery, and behavioral data acquisition were controlled by a Tucker-Davis Technologies iPac system running iCon behavioral interfaces. Auditory stimuli were delivered from a calibrated MF1 speaker positioned approximately 4 cm above the nose poke port, and sound levels were verified with a Larson Davis SoundExpert 821 sound-level meter.

Animals performed a single-interval, two-alternative forced-choice discrimination between 4-Hz and 12-Hz amplitude-modulated (AM) broadband noise (0.1-20 kHz; 100% modulation depth). Gerbils self-initiated a trial by holding in the central nose poke for at least 100 ms, which triggered the acoustic stimulus. Each stimulus had a 100-ms onset ramp followed by a 100-ms unmodulated lead-in before the AM signal. Once the animal left the nose poke region to report its choice, the AM component ended and transitioned into unmodulated noise that remained on during the approach to a food trough. The sound was turned off when the trial ended. Approaching the left trough on 4-Hz trials and approaching the right trough on 12-Hz trials were marked as correct responses and triggered delivery of one 20-mg dustless precision food pellet (Bio-Serv). Incorrect responses were not rewarded and did not produce additional punishment. The two AM rates were presented randomly with equal probability. Stimuli were presented at 55 dB SPL for NH animals and 95 dB SPL for NIHL animals. These levels were selected to compensate for the permanent threshold elevation produced by noise exposure. The resulting sensation levels are quantified in Figure 1F.

### Procedural shaping and acquisition criterion

Before acquisition testing, animals were shaped through the same sequence described previously^21^. During shaping, gerbils learned to approach the food troughs for reward and self-initiate stimulus presentation from the central nose-poke. Shaping was used to establish the procedural requirements of the task before animals entered the two-choice discrimination phase. Once animals reliably initiated trials and approached both troughs, they entered the acquisition phase analyzed in the present study, during which the 4- and 12-Hz stimuli were presented within the same session and animals were required to select the appropriate response on each trial (4-Hz = approach left food trough; 12-Hz = approach right food trough).

Task acquisition was defined a priori as ≥80% correct on both the 4-Hz and 12-Hz stimulus within a single training session. Sessions to criterion was the number of acquisition sessions up to and including the first session meeting that criterion. Trials to criterion was the cumulative number of presented trials over those sessions. Trials were scored as failure when animals did not respond and approach either food trough within a 10 sec response time window. Trials aborted within 100 msec of stimulus presentation were excluded from percent-correct calculations and from the presented-trial count. Aborted trials comprised 496 of 96,763 trials (0.5%).

Training continued until criterion was reached or until performance had ceased to improve, with a maximum of 50 sessions. Three NIHL animals were discontinued without reaching criterion: J_F132 at session 44, and J_M158 and J_M159 at session 50. At discontinuation, none of these 3 animals had come within 5 percentage points of criterion on its poorer AM rate and each had gone at least 11 sessions without a new best performance. For analyses of sessions and trials to criterion, the final training value for these animals was retained as a right-censored lower bound rather than interpreted as a completion time. Sensitivity analyses repeated the acquisition comparison with criteria of 70% and 75% correct on both rates.

### Auditory brainstem response recording

ABRs were recorded before noise exposure and 1, 7, 14, 21, and 28 days afterward. NH animals contributed a single baseline recording. During ABR recording sessions, gerbils were anesthetized with isoflurane (1.0%) and placed in a small acoustic chamber (IAC, Sound Room Solutions). Subdermal pin electrodes were placed at the vertex of the skull (positive), caudal to the left pinna (inverting), and in the left leg (ground). Stimulus generation, presentation, and acquisition were controlled with Tucker-Davis Technologies BioSigRZ software and hardware. Sounds were delivered in closed field through an MF1 speaker coupled to a 10-cm tube positioned at the opening of the left ear canal.

Stimuli consisted of 100-µs clicks and 5-ms tones (2-ms linear ramp) at 1, 2, 4, 8, and 16 kHz. Stimuli were presented from 90 dB SPL downward in 10-dB steps, with 500 repetitions at each level. Threshold was defined as the lowest sound level that elicited a reproducible stimulus-evoked ABR (Figure 1B). When no stimulus-driven response was elicited at 90 dB SPL, the threshold was recorded at the 90 dB SPL ceiling and treated as a lower bound. Ceiling values occurred only for post-noise exposure tone thresholds and not for the day-14 click thresholds used for the primary group comparison.

We performed an exploratory analysis to determine whether noise-induced ABR threshold shifts differed by sex since the NIHL group included both males (n = 7) and females (n = 3). Click-evoked threshold shifts were similar between males and females (44.3 ± 7.9 vs 40 ± 10 dB SPL, respectively; exact permutation test, p = 0.71). Mean threshold shift across tone frequencies did not differ significantly between males and females (32.6 ± 6.1 vs 26.7 ± 12.1 dB SPL, respectively; exact permutation test, p = 0.33). Likewise, no significant sex differences were detected at individual frequencies after correction for multiple comparisons (p>0.05). These results provide no evidence of a sex-dependent difference in the magnitude of NIHL in this cohort.

### Sensation level

Sensation level was calculated for each animal as the sound level used during behavioral training (NH = 55 dB SPL; NIHL = 95 dB SPL) minus that animal’s ABR threshold. NH sensation levels used the animal’s baseline ABR and NIHL sensation levels used the day-14 post-noise exposure ABR. Click sensation level was treated as the primary broadband comparison because the task stimulus was broadband AM noise. Group differences were evaluated with independent-samples t-tests.

### Relating noise-induced hearing loss magnitude to acquisition

For each NIHL animal, threshold shift was calculated as the day-14 ABR threshold minus the pre-noise exposure baseline for clicks and for each tone frequency. A mean tone threshold shift was obtained by averaging across 1, 2, 4, 8, and 16 kHz. Click threshold shift and mean tone shift were compared with sessions and trials to criterion using Spearman rank correlations, producing four comparisons that were Holm-corrected. Spearman correlations were used because the sample per comparison group was small (n = 10), thresholds were quantized in 10-dB steps, and three acquisition outcomes were right-censored to lower bounds. Absolute day-14 thresholds and shifts at individual tone frequencies were also examined.

### Measures of task engagement

Four task engagement measures were computed for each animal. Trials per session was the mean number of presented trials completed per training session. Trial failure rate was the proportion of presented trials on which the animal did not make a choice within the response time window. Abort rate was the proportion of all logged trials on which the animal left the nose poke region within the first 100 msec of stimulus presentation, which was prior to AM onset. Response latency was measured from AM onset to the animal’s departure from the nose poke region on trials in which a choice was made. Latency distributions were typically right-skewed, and thus each animal was summarized by its median latency. Group comparisons used one value per animal and nonparametric Mann-Whitney U tests.

### Video recordings

Behavioral sessions were recorded from above with a USB camera at 60 frames per second (Basler ace2) and multiple body part positions (nose, head, ears, back, and base of tail) were tracked frame by frame with SLEAP^52^. Tracked points with SLEAP confidence scores <0.3 were set to NaN and excluded from subsequent analyses. We generated a trained gerbil network that consisted of >11,000 iterations and labeled frames. Video recordings were available for 14 animals (NH, n = 8; NIHL, n = 6) because the video-recording system was implemented after behavioral data collection had begun for six animals. The tracked arena extent was defined dynamically from the 0.5^th^ to 99.5^th^ percentiles of tracked positions across all body parts to reduce the influence of occasional tracking outliers. Median positions of the nose poke and two food troughs were used to define the task landmarks. For each task session, the region of interest was defined from these landmarks, expanded by 5% of the dynamic arena extent, and clipped to the tracked arena boundary. Head positions were binned into 5-pixel squares, and percentage of arena occupied was calculated as the proportion of bins visited at least once.

The percentage of arena occupied increases mechanically with the number of trials in a session because longer sessions provide more opportunities to visit a bin. Thus, primary analyses were restricted to sessions containing ≥100 presented trials. The change across training was quantified within each animal by the Spearman correlation between session number and percentage of arena occupied and per-animal coefficients were tested against zero with Wilcoxon signed-rank tests. We also compared the first and second half of each animal’s training. Between-group comparisons used nonparametric Mann-Whitney U tests on one value per animal.

For each correct trial, the tracked head trajectory was extracted from the nose poke to the selected food trough. Trajectories containing fewer than 10 valid tracked frames were excluded. Each remaining trajectory was resampled to 100 equally spaced points along arc length so that the measure reflected route geometry rather than running speed. Leftward and rightward trials were analyzed separately. A path-spread estimate required at least three valid trajectories. Within each session, trajectories ending at the same trough were averaged point-by-point to obtain a mean route. Path spread was calculated as the mean Euclidean distance of individual trajectories from the corresponding session mean route across the 100 resampled points and was then averaged across the two troughs.

Primary path-spread analyses used the same ≥100 trial/session criterion as the percentage of arena occupied analysis so that both kinematic measures described the same sessions. Change across training was quantified by a per-animal Spearman correlation between session number and path spread. Coefficients were tested against zero within each group with Wilcoxon signed-rank tests and compared between groups with a nonparametric Mann-Whitney U test. The between-group result was also evaluated using the change from the first to last quarter of normalized training sessions, the change from the first to second half of training sessions, and normalized progress in place of session number.

### Per-session logistic regression model

To quantify the information influencing behavioral choice, each training session was fit separately with a logistic regression:

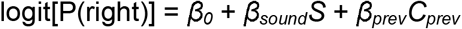

where *S* = +1 for 12-Hz trials and −1 for 4-Hz trials, *C_prev_*= +1 when the previous choice was right and −1 when it was left, β*_0_* is the side-bias/intercept term, β*_sound_*is weight of the current sound, and β*_prev_* is the weight of the previous choice. The sound weight (β*_sound_*) therefore quantified how strongly stimulus identity influenced the animal’s current choice, whereas the side-bias and previous-choice terms captured choice tendencies that were independent of the current acoustic stimulus. Only trials on which the animal made a left or right choice (i.e., correct and incorrect trials) entered the model. The first trial of each session was excluded because no previous choice was available, and sessions with fewer than 40 scored trials were not fit. This excluded 12 sessions and yielded 437 total fitted sessions. A ridge penalty (λ = 1) was applied to the two slope terms, with the intercept left unpenalized, to prevent divergence in sessions with complete or near-complete separation. Side bias and previous-choice weights were analyzed in absolute value when quantifying the magnitude of sound-independent choice tendencies.

Within-animal comparisons across training was summarized by the Spearman correlation between session number and each fitted weight. For between-group analyses, each animal’s training record was divided into four quarters of normalized training progress and reduced to one mean value per quarter. Nested random-intercept linear mixed-effects models were fit by maximum likelihood with animal as the grouping factor, were compared by likelihood ratio. Adding hearing status to a model containing quarter tested the group difference in level and adding the hearing status x quarter interaction tested whether the shape of the trajectory differed between groups. Quarter-wise post hoc comparisons used nonparametric Mann-Whitney U tests with Holm correction.

### Trial-by-trial logistic regression model

The same three choice influences of sound, side bias, and previous choice, were also fit with a dynamic logistic model in which the weights could evolve continuously across trials^22^. The observation model was identical to the per-session logistic regression model, but each weight was allowed to change from one trial to the next according to a Gaussian random walk:

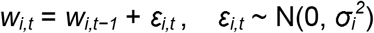

where *i* indexes the sound, side-bias, or previous-choice weight, and *t* indexes the trial number. The step-size parameter σ*_i_* was estimated from the data for each weight, with a separate step size across session boundaries. Hyperparameters were fit by maximizing model evidence, so the smoothness of each trajectory was estimated rather than imposed. Models were fit with PsyTrack^22^ with one model per animal across their complete training record. All choice trials were included, yielding 94,871 trials in 449 sessions.

Since adjacent trial estimates from a smoothed random walk are highly correlated, all inferential tests were performed on one summary value per animal rather than treating trials as independent observations. The start value was the mean weight over the first 200 trials and the end value was the mean over the final training session. Within-group comparisons were tested with Wilcoxon signed-rank tests on each animal’s start-to-end difference and between-group comparisons used nonparametric Mann-Whitney U tests. For normalized training progress analyses, the primary summary was the area under each animal’s training trajectory from 0 to 1 progress. Linear mixed-effects models on quarter means were then used to test effects of training progress, hearing status, and their interaction.

For this two AM rate discrimination task, the model would support 80% correct on both AM rates when the sound weight exceeded ln(4) plus the absolute side-preference weight, where ln(4) ≈ 1.39 corresponds to the log-odds of an 80% choice probability. This level is shown in Figure 6C-E only as a reference. The primary acquisition endpoint throughout the study remains the observed session-level criterion of ≥80% correct on both rates.

## Acknowledgements

This work is supported by grants to JDY (NIH NIDCD R01DC023926 and the Busch Biomedical Grant Program from Rutgers University).

## Author contributions

MC, MM, and WM performed the experiments; MC, MM, WM, SA, KPARN, and JDY analyzed the data; TMM and JDY designed the experiments; MC, MM, TMM, and JDY wrote the paper.

## Data availability statement

Data and analysis code can be found at https://rutgers.box.com/v/Auditory-Learning-Data.

---

**Supplementary Figure S1.**
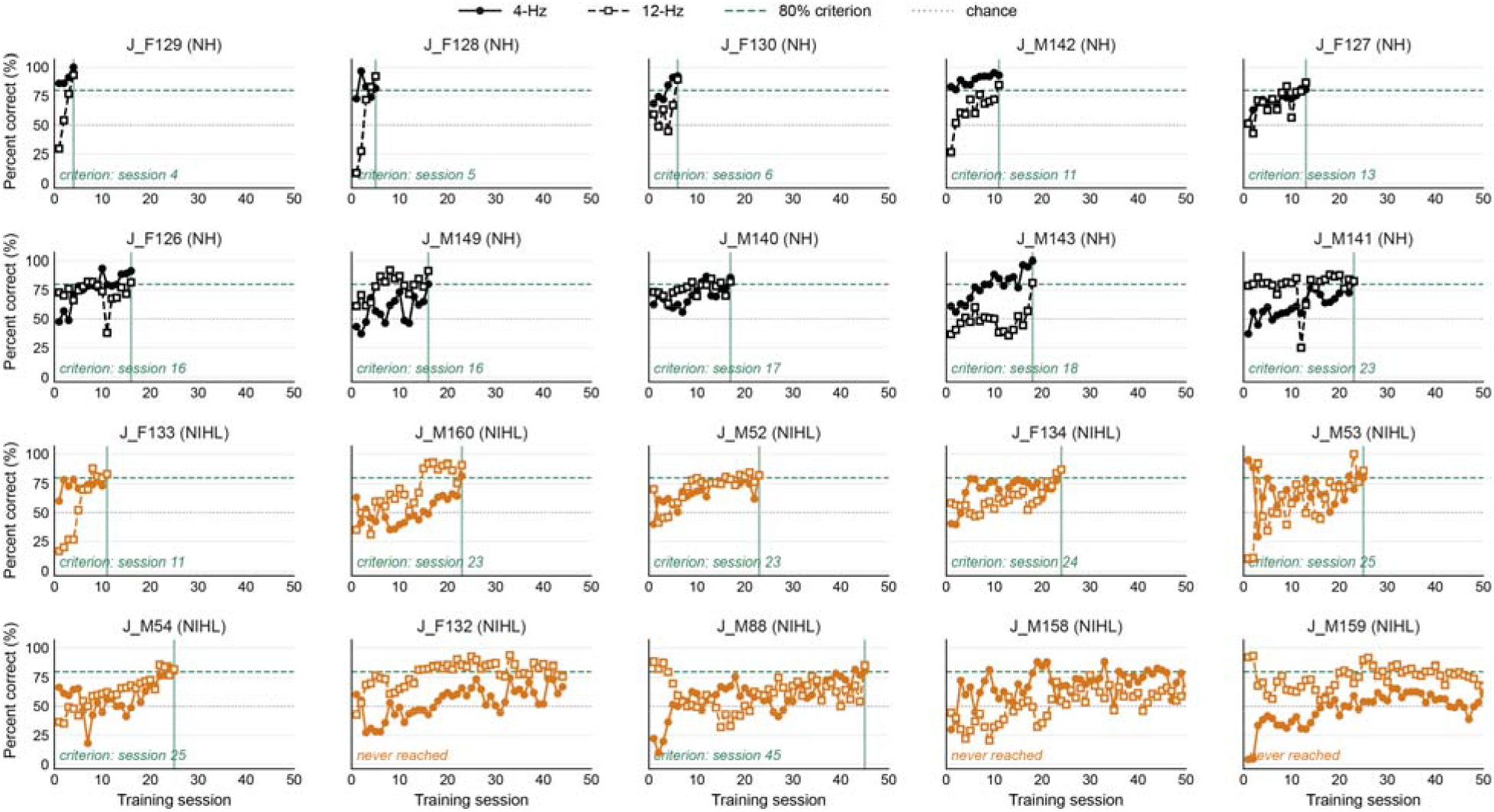
Learning curves across training sessions for all animals. Percent correct for the 4-Hz (solid line, filled circles) and 12-Hz (dashed line, open squares) AM rates is shown for all normal-hearing (black) and NIHL (orange) animals. Animals are ordered within each hearing status group by the number of sessions required to reach criterion. The horizontal green dashed line marks the 80% acquisition criterion and the grey dotted line marks chance performance (50%). For animals that reached criterion, the vertical green line indicates the first session in which performance was ≥80% correct for both AM rates. J_F132, J_M158, and J_M159 were discontinued without reaching criterion.

**Supplementary Figure S2.**
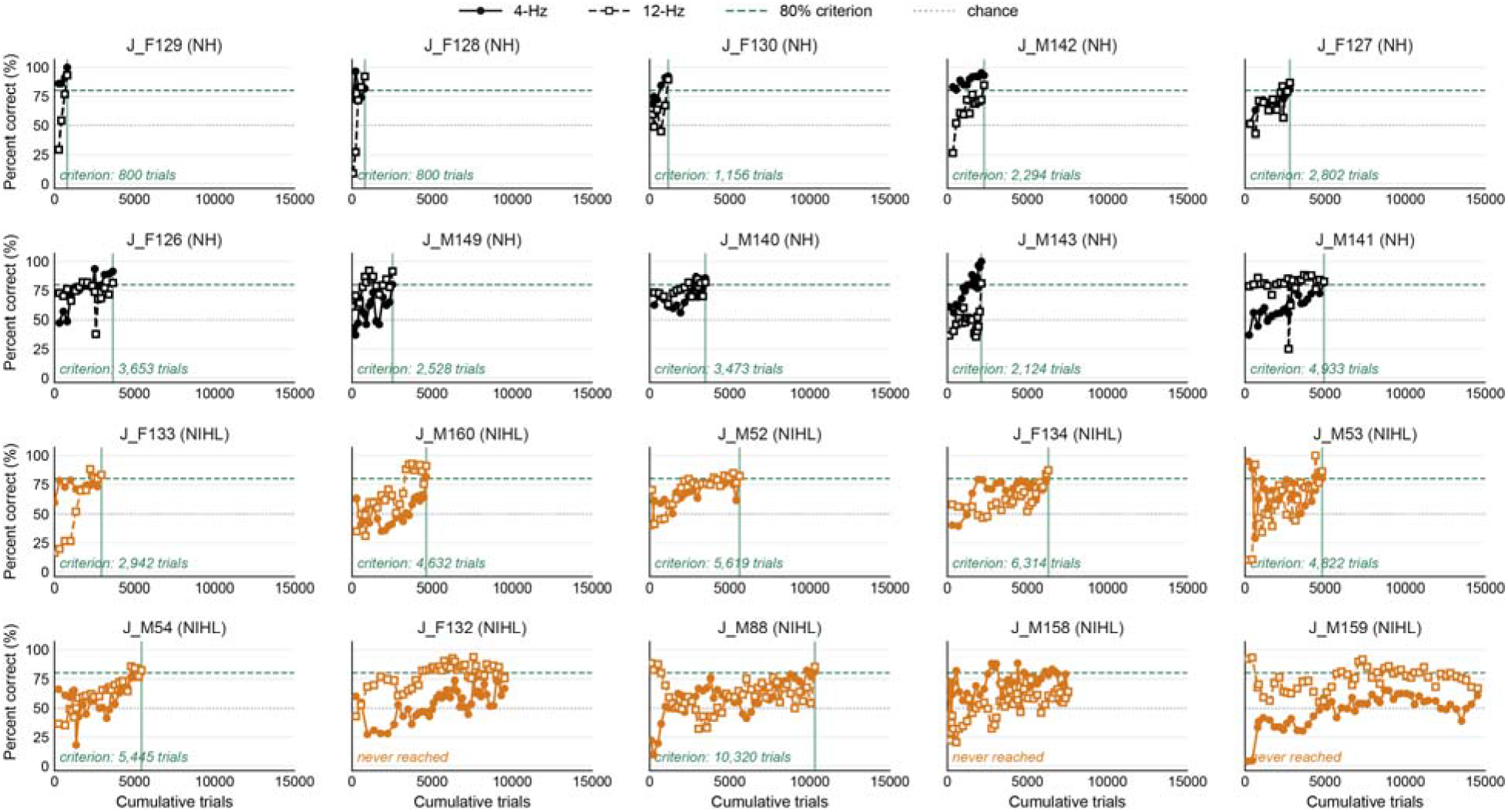
Learning curves across cumulative trials for all animals. Percent correct for the 4-Hz (solid line, filled circles) and 12-Hz (dashed line, open squares) AM rates is shown for all normal-hearing (black) and NIHL (orange) animals. Animals are ordered within each hearing status group by the number of sessions required to reach criterion, matching Supplementary Figure 1. The horizontal green dashed line marks the 80% acquisition criterion and the grey dotted line marks chance performance (50%). For animals that reached criterion, the vertical green line indicates the cumulative trial count at the first session in which performance was ≥80% correct for both AM rates. J_F132, J_M158, and J_M159 were discontinued without reaching criterion. N = 10 animals per hearing status group.

**Supplementary Figure S3.**
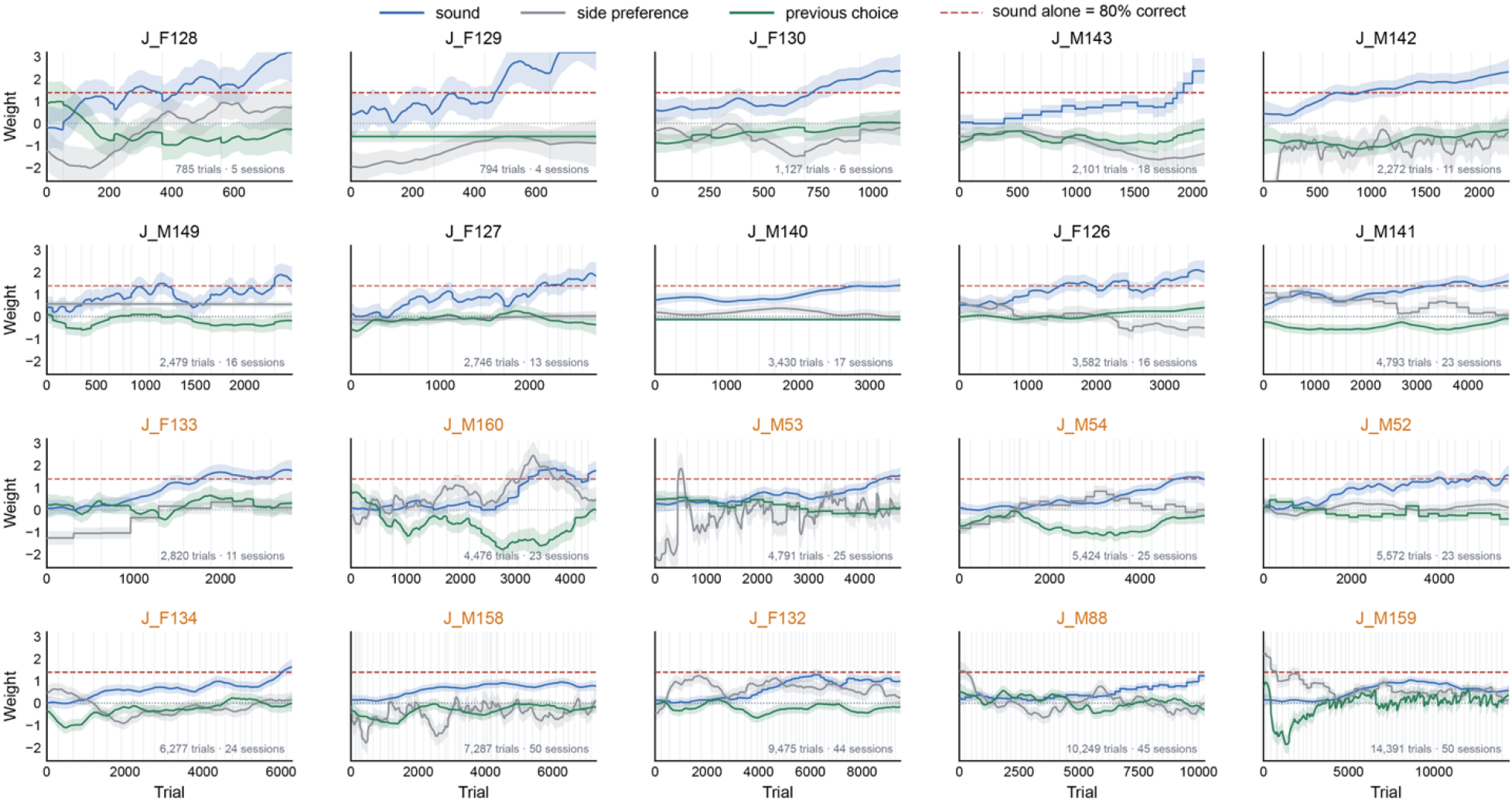
Trial-by-trial dynamic logistic regression fits for all animals. Trial-by-trial model weights are shown across the complete training record for all NH (black animal labels) and NIHL animals. Blue lines represent sound weight, gray lines represent side-bias weight, and green lines represent previous-choice weight. Shaded regions indicate ± 2 posterior SD. Vertical gray lines indicate session boundaries. The horizontal red dashed line marks the sound weight at which the sound alone would support 80% correct performance. The total number of trials and training sessions included in each fit is indicated within each panel. Animals are ordered by hearing status group and total number of training trials.

